# Multi-agent AI enables evidence-based cell annotation in single-cell transcriptomics

**DOI:** 10.1101/2025.11.06.686964

**Authors:** Gautam Ahuja, Alex Antill, Yi Su, Giovanni Marco Dall’Olio, Sukhitha Basnayake, Göran Karlsson, Parashar Dhapola

## Abstract

Cell type annotation remains a critical bottleneck, with current methods often inaccurate and requiring extensive manual validation, particularly in disease contexts. While large language models (LLMs) show promise, they can be unreliable due to hallucinations. We developed CyteType, a multi-agent framework that generates competing hypotheses grounded in full expression data and study context, validates against external databases, and iteratively self-evaluates. Comprehensive benchmarking demonstrates that CyteType substantially outperforms reference-based and LLM-based methods, with self-generated confidence scores reliably identifying trustworthy annotations. CyteType transforms cell type annotation from label assignment into evidence-grounded biological discovery.

Python (AnnData compatible): https://github.com/NygenAnalytics/CyteType

R (Seurat compatible): https://github.com/NygenAnalytics/CyteTypeR

## Main text

Single-cell RNA sequencing datasets now routinely contain millions of cells ^1,2^. As computational methods are becoming increasingly more efficient, the bottleneck has shifted from data analysis to biological interpretation ^3,4^, particularly in identifying cell type identities of clusters. Cell type annotation remains most challenging in disease contexts ^5,6^, and current reference-based classifiers trained on healthy tissue exhibit an accuracy drop of 15-30% when applied to disease samples ^7^, missing rare cell types in approximately 20% of cases. Manual annotation can lead to superior results over automated methods but is time-consuming and shows 25% inter-annotator variability ^8^. Additionally, current methods provide only cell type labels without justification or caveats ^9^, limiting result interpretability. Large language models (LLMs) have shown potential for cell type annotation, with GPT-4 achieving 75% agreement with expert annotations ^10^. However, existing LLM approaches have significant limitations: they process only top marker genes rather than full expression profiles, rely on static knowledge from pretraining data ^11^, and lack mechanisms to validate predictions against external databases or quantify uncertainty.

We developed CyteType, a multi-agent framework that addresses these limitations through three key innovations (Figure 1A). First, CyteType implements hypothesis-driven reasoning, systematically testing competing cell type interpretations against full expression profiles, pathway enrichment, and study context to mimic expert annotation workflows. Second, it validates predictions by simulating an expert panel that performs automated reference checking against marker databases ^12–14^. Third, it provides comprehensive biological context by connecting annotations to current literature, disease associations, and therapeutic targets. This framework transforms cell type annotation from a classification task into an evidence-based characterization process with interpretable reasoning and testable hypotheses.

**Figure 1.**
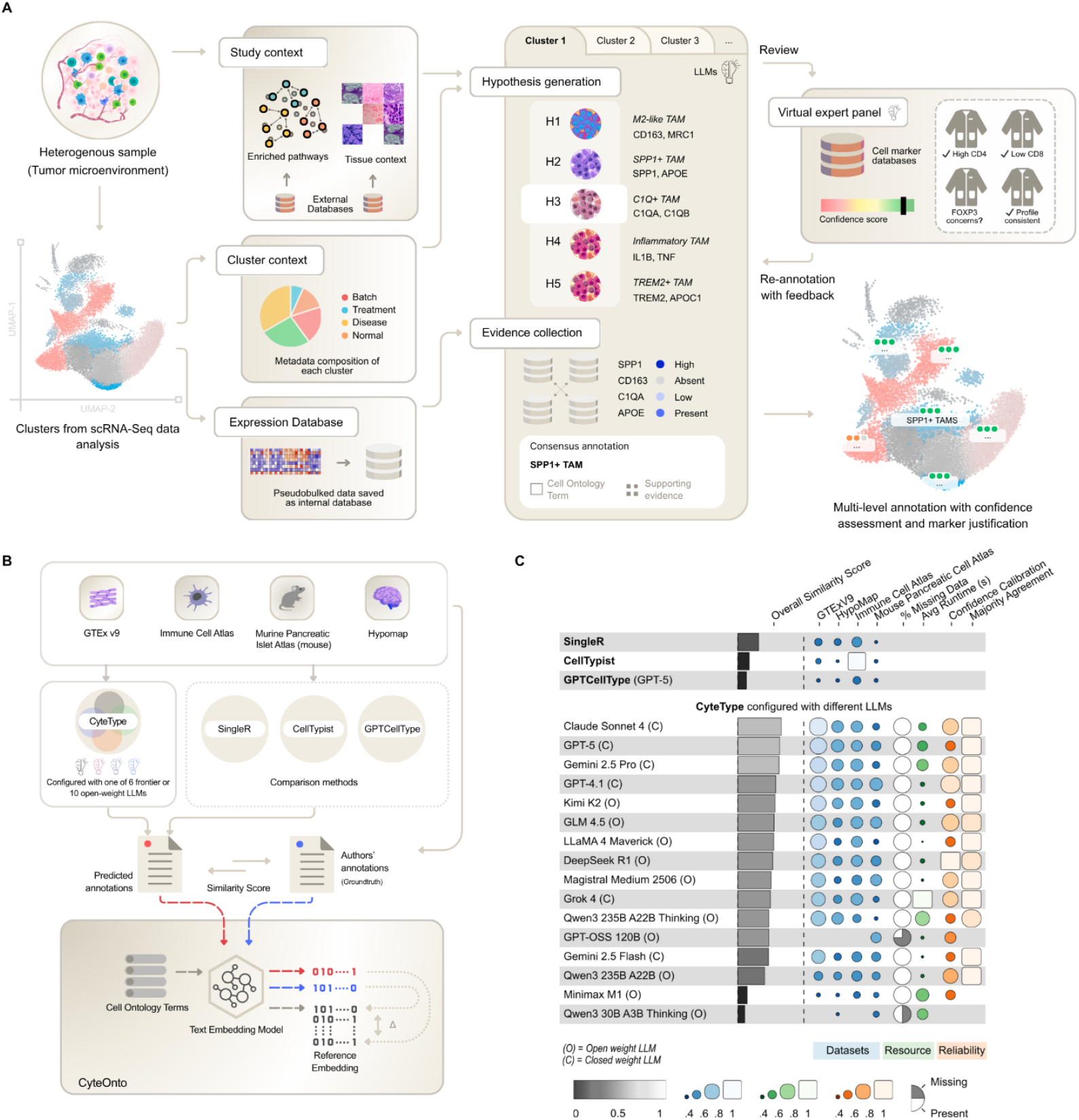
CyteType overview and benchmarking. (A) Workflow schematic of CyteType for automated cell type annotation from scRNA-seq clusters. Study and cluster contexts are built from pathway analysis and metadata. LLM agents generate cell type hypotheses and collect supporting evidence by querying a pseudo bulked expression database. The top hypothesis is selected based on evidence strength and linked to Cell Ontology terms. A reviewer agent establishes a virtual expert panel that assigns confidence scores and triggers re-annotation if needed. (B) Benchmarking strategy comparing CyteType against three existing methods across four diverse tissue datasets. Prediction accuracy was assessed using CyteOnto, which calculates similarity score between predicted and author-provided annotations in the Cell Ontology embedding space. (C) Benchmarking results showing overall and dataset-specific similarity scores, along with reliability and performance metrics across multiple LLM models.

Rigorous benchmarking of cell type prediction algorithms has proven challenging due to the free-text nature of cell type labels. To address this, we developed CyteOnto, a semantic similarity framework that maps cell type labels to Cell Ontology terms using text embeddings of LLM-augmented descriptions (Figure 1B). Unlike string-matching or graph-based methods that fail to capture semantic relationships, CyteOnto maps predicted and ground-truth (authors annotation) to their nearest Cell Ontology terms in embedding space, then quantifies normalized distances between these terms to derive a kernel-induced similarity score ranging from 0 to 1. Validation against curated test cases demonstrated that CyteOnto preserves biological meaning, maintains monotonic similarity score decay across cell type hierarchies, and outperforms traditional ontology distance metrics (Supplementary Information 1). This framework enabled quantitative comparison of CyteType against existing methods using similarity scores rather than exact or fuzzy label matching.

We evaluated CyteType on four benchmark datasets ^15–18^ spanning 205 clusters, across diverse biological contexts (Figure 1B), using author provided annotations as ground truth proxy. To measure accuracy of predicted cell types from each benchmarked methods, we used CyteOnto similarity scores against the ground truth proxies. To isolate architectural benefits of CyteType from underlying LLM capabilities, we compared CyteType against GPTCellType ^10^ using identical LLM (GPT-5), alongside established reference-based methods: CellTypist ^18^ and SingleR ^19^. Using identical models (GPT-5), CyteType significantly outperformed GPTCellType with 388.52% higher similarity score across four datasets (*z* = 8.04, *p* < .001, Tukey-adjusted), demonstrating that structured hypothesis testing and evidence collection improve upon direct LLM prompting (Figure 1C). This performance-gain demonstrates that CyteType’s architectural innovations provide value independent of underlying LLM capabilities. CyteType’s median similarity score also exceeded those from reference-based methods, CellTypist (*z* = 8.15, *p* < .001) and SingleR (*z* = 8.36, *p* < .001), with performance gains consistent across diverse biological contexts but most pronounced in multi tissue datasets. CyteType showed 267.9% and 100.67% higher average similarity score over CellTypist and SingleR, respectively. One notable exception was CellTypist’s near-perfect (similarity score: median: 1.0, mean: 0.8) performance on the immune cell atlas dataset, where author labels were generated using CellTypist itself, serving as a positive control for CyteOnto’s similarity scoring. Hence, CyteType outperformed existing cell annotation methods by an average of 252.36% higher similarity score. To determine whether CyteType’s strong performance arose from the underlying language model or from CyteType’s framework and structured reasoning, we evaluate its robustness across diverse LLMs.

To assess robustness of CyteType’s architecture, we benchmarked 16 LLMs spanning closed-weight (e.g. GPT-5, Claude Sonnet 4, Gemini 2.5 Pro) and open-weight models (DeepSeek R1 ^21^, Qwen3 Thinking) (Figure 1C). Closed-weight models achieved the highest mean similarity score, while open-weight models showed modest but significantly decreased (*b* = -0.035, *SE* = 0.011, *p* < .001) mean score yet still outperformed traditional methods. Notably, LLMs with built-in chain-of-thought reasoning demonstrated no significant advantage (*b* = 0.014, *SE* = 0.011, *t*(3977), *p* = 0.22) over standard models, suggesting CyteType’s structured workflow supersedes model-native reasoning capabilities. This finding has practical implications: researchers can select models based on cost and privacy requirements without sacrificing the benefits of CyteType’s architecture. Closed-weight models (GPT-5, Claude Sonnet 4) maximize accuracy for critical applications, while open-weight models (Kimi K2 ^20^, Deepseek R1) achieve 95% of peak performance at low cost, suitable for exploratory analysis or budget-constrained settings (Supplementary Information 2).

Having established CyteType’s performance and robustness across LLMs, we next evaluated whether its predictions can be trusted and systematically interpreted. CyteType’s Reviewer agent (Figure 1A) generates calibrated confidence scores and heterogeneity flags that enable targeted expert review. High-confidence annotations were significantly more likely to have higher similarity score than those with low-confidence (*F* = 23.88, *p* < .001), while heterogeneous clusters showed significantly lower similarity score against author labels (*F* = 8.45, *p* < .01). Across repeated runs, median majority agreement exceeded 80% for all LLMs, with consensus Cell Ontology labels achieved for >70% of clusters. Both reviewer confidence and heterogeneity flags predicted similarity scores even after accounting for LLM choice (likelihood ratio test: *χ*^2^ = 42.06, *df* = 16, *p* < .001 and *χ*^2^ = 35.44, *df* = 16, *p* < .01, respectively). These metrics create an interpretable “trust layer” that existing annotation methods lack, transforming cell type annotation from a black-box classification into a transparent process with quantified uncertainty.

Finally, we evaluated whether CyteType’s evidence-based framework extends beyond accurate classification to enable refinement of existing data potentially leading to novel discoveries. Applying CyteType to 977 clusters across 20 datasets revealed systematic patterns. Rather than simply validating existing annotations, CyteType added value across all scenarios: 41% of clusters received functional enhancement (e.g., adding cell state information), 29% underwent refinement to specific subtypes, and 30% required major reannotation (Supplementary Information 3). Importantly, this distribution demonstrates that CyteType functions not merely as a classifier but as a discovery tool, uncovering biological context missed by reference-based methods. Annotations mapped to 327 unique Cell Ontology terms with high diversity (no term exceeding 2.5% frequency) and identified 116 distinct cell states, with activation (37%) and maturation (16%) most prevalent. Predicted heterogeneity strongly correlated with lower confidence (Pearson’s correlation coefficient: *r* = -0.79, *p* < .001), validating the system’s ability to flag mixed populations. High-confidence discrepancies occurred predominantly in disease contexts where reference methods historically underperform, suggesting CyteType uncovers biologically meaningful refinements. For example, in the diabetic kidney disease atlas author denominated parietal epithelial cells and leukocytes were relabelled injured proximal tubule cell (ALDH1A2+, CFH+, VCAM1+) and T cell (activated CD45+DOCK2+ pro-inflammatory) respectively.

CyteType is available as Python (AnnData ^22^ compatible) and R packages (CyteTypeR for Seurat ^23^ compatibility) and generates comprehensive reports that integrate directly into existing workflows. Cell type annotations are returned into AnnData/Seurat objects along with Cell Ontology terms for downstream analysis and data integration. Interactive HTML reports provide detailed marker justification, pathway enrichment, literature evidence, and disease associations, with embedded chat functionality enabling iterative refinement and hypothesis exploration.

CyteType addresses fundamental limitations in current annotation approaches by transforming cell type identification from classification into evidence-based characterization. While reference-based methods constrain predictions to previously characterized cell types and degrade under domain shift, and transformer-based foundation models like scGPT ^24^, Geneformer ^25^ and TranscriptFormer ^26^ require careful fine-tuning for novel contexts, CyteType’s hypothesis-testing framework discovers cell states and rare populations with calibrated uncertainty. Recent LLM approaches like GPTCellType ^10^ and Cell2Sentence ^13^ have demonstrated that generative models can produce interpretable annotations; however, our direct comparison using identical models shows that multi-agent architecture provides substantial value beyond the capabilities of LLM alone. CyteType’s built-in quality control, through confidence scores and heterogeneity detection, enables computational biologists to efficiently triage results, accepting high-confidence annotations while directing expert attention to uncertain clusters. By connecting annotations to pathway enrichment, literature, and disease contexts, CyteType delivers not just labels but biological insight with testable hypotheses, closing the gap between automated annotation and biological understanding.

## Software and Code Availability

CyteType SDKs are available on Github:

Python (AnnData compatible): https://github.com/NygenAnalytics/CyteType

R (Seurat compatible): https://github.com/NygenAnalytics/CyteTypeR

CyteOnto is available as an open-source Python package at https://github.com/NygenAnalytics/CyteOnto. Analysis code, benchmark datasets, and processed results are available at https://github.com/dalloliogm/cytetype_benchmark/tree/main/notebooks/manual and https://github.com/NygenAnalytics/CyteType_manuscript. Interactive HTML reports with embedded chat functionality are generated automatically for each dataset. An example HTML report for Human Tumour Atlas Network (HTAN) MSK can be browsed at https://nygen-labs-prod--cytetype-api.modal.run/report/5b4eb3e1-fde7-4609-8be0-2bea015c241d?v=250722

## METHODS

### CyteType Architecture and Implementation

CyteType is a multi-agent system designed to automate the rigorous scientific reasoning process for **cell type annotation**. The CyteType architecture breaks the annotation task into five distinct, collaborative agents. The **Contextualizer** agent sets up the context by inferring organism, tissue, and pathway context before annotation and establishing biological ground truth. The **Annotator** agent employs a hypothesis-driven framework, generating and systematically testing multiple biological interpretations for each cell cluster to facilitate evidence-based identification. The **Reviewer** agent provides a critical validation, using a simulated panel of domain experts and external reference databases to assess annotation confidence and detect cellular heterogeneity. The **Literature & Clinical Relevance** agent connects the findings to PubMed literature, Disease Ontology, and Drug Ontology. Finally, the **Summarizer** agent synthesizes the results into a coherent, study-wide biological narrative, which can be viewed as a report on a browser. Additionally, the **Chat** agent and user-triggered reannotations with feedback enable users to introspect agentic reasoning, query data, and update annotations.

**Table 1:**
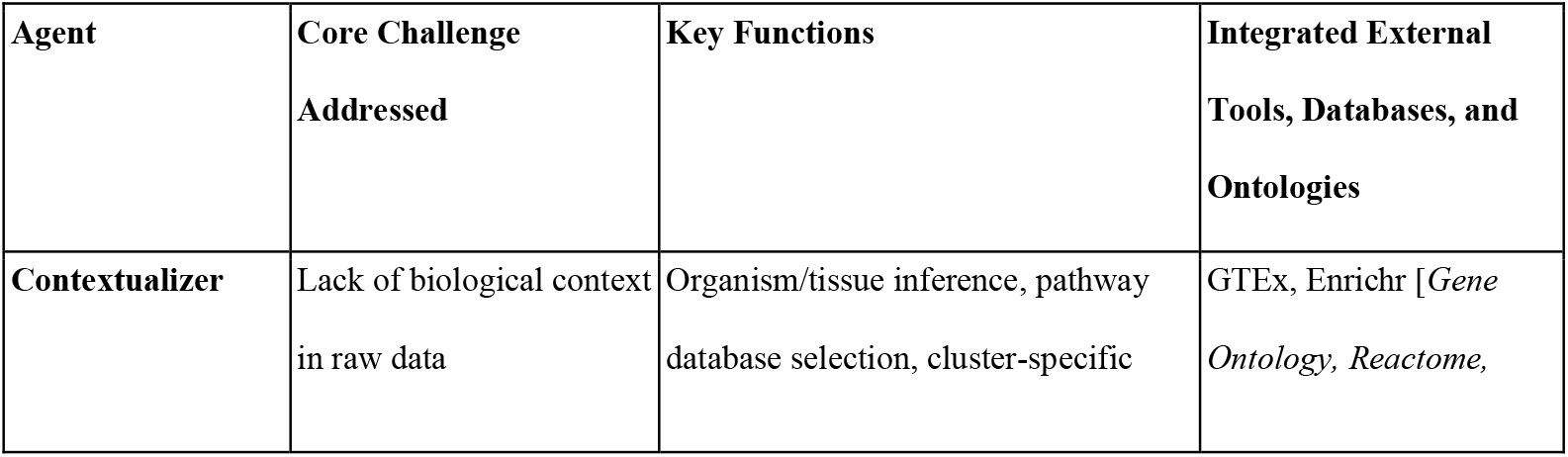
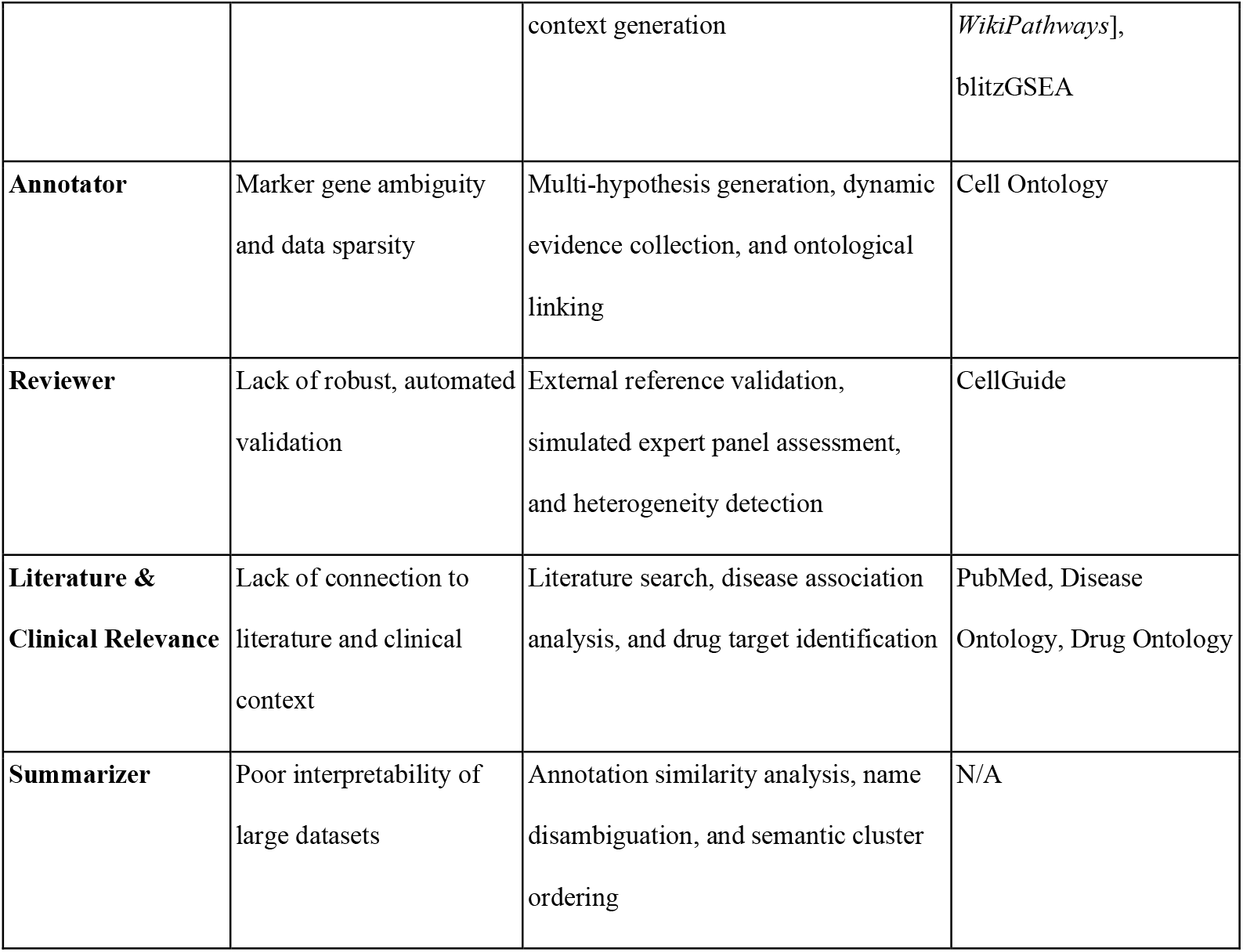
Overview of the CyteType Multi-Agent Architecture.

CyteType is designed as a resilient and scalable API-driven service that integrates multiple specialized AI agents, external biological databases, and large language models (LLMs) to perform end-to-end cell type annotation.

### Data Ingestion and Preparation

The workflow begins with data ingestion with standard single-cell data formats like AnnData objects (.h5ad) and Seurat (.RDS), as well as direct data submission via API calls. Along with the expression data, users provide essential study context (e.g., species, tissue source, experimental design). This information is consolidated into an internal query-ready database that stores the expression percentages for each gene in each cluster, top marker genes for each cluster, and associated percentages across metadata categories for efficient access by the agentic system.

### Integration with LLMs and External Databases

The agents are powered by a suite of LLMs (e.g., GPT-5, Claude Sonnet 4, Gemini 2.5 Pro, etc.) equipped with function-calling abilities. This allows the agents to interact programmatically with a wide array of external biological databases, which are available as callable “tools.” These resources include ontologies (Cell Ontology, Gene Ontology), literature databases (PubMed), and specialized molecular databases (GTEx, CellGuide, Drug Ontology), ensuring that all reasoning is grounded in established scientific knowledge. Tool usage budgets are enforced per agent with automatic prompt constraints when approaching limits.

### System Outputs and Integration

CyteType delivers its results through multiple formats to suit different user needs. A structured JSON output provides granular annotations and ontology IDs for computational use. A comprehensive HTML report offers a user-friendly view of the results, including an interface for human-in-the-loop feedback and re-annotation via a chat portal, enabling interactive, human-led investigation of the data. The final annotations can be integrated back into the original AnnData or Seurat objects, allowing researchers to proceed with familiar downstream analysis and visualization tools. This entire workflow is deployed on a scalable cloud infrastructure, enabling the parallel processing of hundreds of clusters simultaneously and making it suitable for large-scale single-cell transcriptomic studies.

### Scalable Processing

The system implements concurrent cluster processing with configurable parallelism (default 20 clusters, scalable to hundreds simultaneously). Processing time averages 5-10 minutes per cluster but can vary depending on speed of chosen LLM and model provider’s inference speed, and the number of self-iterations, consuming between 400,000 and 600,000 tokens per cluster. Pathway enrichment analysis executes via distributed compute functions with horizontal scaling and isolation.

### LLM Provider Management

CyteType employs multi-provider rotation with primary/secondary/fallback selection and preflight validation. Agent execution implements unified retry policies with exponential backoff (5s → 10s → 30±30s → 60±60s) for HTTP and model output structure validation errors. The system supports deployment via cloud APIs (OpenRouter, AWS Bedrock) or locally hosted Ollama models with configurable timeout and scaling parameters. Per-call cycling ensures diversity and resilience while usage limits prevent cost overruns.

## Benchmark Datasets and Evaluation

### Dataset Selection

Four benchmark datasets were selected to span diverse biological contexts: HypoMap (developing mouse brain, 66 clusters), Immune Cell Atlas (human immune cells, 45 clusters), GTEx v9 (cross-tissue human atlas, 74 clusters), and Mouse Pancreatic Cell Atlas (pancreatic tissue, 20 clusters). Datasets were down sampled to reduce computational requirements while preserving cellular diversity (Table SI-3-1, with down sampling notebooks available at https://github.com/dalloliogm/cytetype_benchmark/tree/main/notebooks/manual and https://github.com/NygenAnalytics/CyteType_manuscript).

### Comparison Methods

CyteType was benchmarked against three established methods representing different annotation paradigms:

1. *CellTypist* (Python interface v1.7.1): Reference-based supervised logistic regression classifier. Pre-computed models were selected per dataset: Developing_Mouse_Brain (HypoMap), Immune_All_High (Immune Cell Atlas), Adult_Human_PancreaticIslet (Mouse Pancreatic Cell Atlas), and tissue-specific models (GTEx v9).
2. *SingleR* (R Bioconductor): Reference-based correlation method using SingleR. Reference datasets matched testing contexts: HumanPrimaryCellAtlasData (Immune Cell Atlas and GTEx v9) and MouseRNAseqData (Mouse Pancreatic Cell Atlas and HypoMap).
3. *GPTCellType*: LLM-based annotation using differential gene expression. Implemented in R following published methods, utilizing Seurat’s FindMarkers() framework for marker identification, followed by GPT-5 queries via OpenAI R wrapper (openai v0.4.1).
4. All comparison methods were run following published user guidelines. Code and processed outputs are available at https://github.com/dalloliogm/cytetype_benchmark/tree/main/notebooks/manual and https://github.com/NygenAnalytics/CyteType_manuscript

### Multi-LLM Benchmark

Sixteen LLMs were evaluated, meeting selection criteria: context window >32,000 tokens, input cost < $ 5 per million tokens, accessible via OpenRouter, tool-use capabilities, and Graduate-level Google-proof Q&A Benchmark score >65. Models were run via OpenRouter on the lowest-cost 8-bit floating point provider when available (as of August 29, 2025). LLMs were excluded if superseded by newer versions.

### Large-Scale Analysis

CyteType was applied to 20 datasets comprising 977 clusters (Supplementary Information 3). Annotations were compared against original author labels using CyteOnto semantic similarity, categorized as high (66.6-100%), moderate (33.3-66.6%), or low (0-33.3%) concordance. Cell states and Cell Ontology term diversity were analysed across all predictions.

### CyteOnto Semantic Similarity Framework

#### Similarity Computation

CyteType annotations were mapped to the nearest Cell Ontology term based on cosine similarity between generated description embeddings and precomputed Cell Ontology embeddings. Semantic similarity between two annotations was computed using a Gaussian Hill Kernel over cosine similarity (GHKcos) with optimized parameters (σ = 0.25, center = 1.0, amplitude = 1.0):

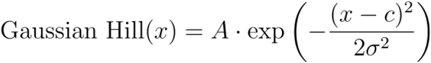

where *c* is the center (default 1.0),σ controls the width (optimally 0.25),*A* is the amplitude (default 1.0), and *x* is the raw cosine similarity score. This transformation emphasizes high-similarity relationships while maintaining sensitivity across the similarity spectrum, providing smooth monotonic decay across biological gradients (detailed methodology in Supplementary Information 1).

#### Validation

CyteOnto was validated using three curated test datasets: (1) 30 Cell Ontology term pairs with defined biological relationships, (2) CD4+ T cell reference series with 40 progressively distant cell types, and (3) sensory neuron reference series. The GHKcos metric demonstrated superior performance over string-matching, graph-based, and direct embedding approaches in preserving biological meaning and maintaining monotonicity (Supplementary Information 1).

### Statistical Analysis

#### Linear Mixed Effects Models

##### Majority Agreement Analysis

The impact of LLMs on categorical cell ontology prediction across multiple datasets and iterations was examined via a generalised linear mixed effects model.

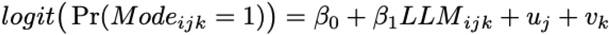

Where *β*_0_ is the overall intercept, *β*_1_ represents the fixed effect of LLM, 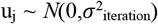 is the random intercept for interaction *j*, 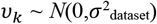 is the random intercept for dataset *k*, and the residual variance follows a binomial distribution. Model parameters were estimated with maximum likelihood with Laplace approximation.

### Similarity to Author Analysis

The impact of LLMs on similarity to author predictions across multiple datasets was examined via an LME.

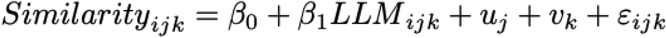

Where *β*_0_ is the overall intercept, *β*_1_ represents the fixed effect of LLM, 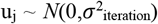 is the random intercept for iteration *j*, 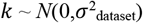 is the random intercept for dataset *k*, and *ε*_*ijk*_ represents the residual error. Model parameters were estimated with restricted maximum likelihood.

### Heterogeneity Analysis

To monitor the performance of LLM’s heterogeneity ratings across multiple datasets, an LME was devised.

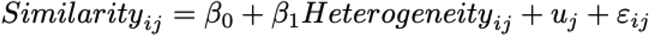

Where *β*_0_ is the overall intercept, *β*_1_ represents the fixed effect of LLM, 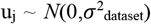 is the random intercept for dataset *j*, and 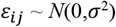 is the residual error for observation *i* in dataset *j*. Model parameters were estimated with restricted maximum likelihood.

An F1 score was derived to measure the ability of LLMs to predict inaccurate cell type predictions via heterogeneous clusters prediction. Similarity to the author score was dichotomised by the midpoint between two centroids from k-means clustering (*k* = 2). The effect of LLM’s performance across multiple datasets was examined via an LME.

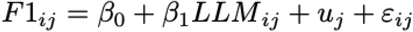

Where *β*_0_ is the overall intercept, *β*_1_ represents the fixed effect of LLM, 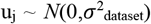 is the random intercept for dataset *j*, and *ε*_*ij*_ ∼ *N*(0,*σ*^2^) is the residual error for observation *i* in dataset *j*. Model parameters were estimated with restricted maximum likelihood (REML)

### Confidence Analysis

To monitor if confidence predicted the similarity to the author score, an LME was used.

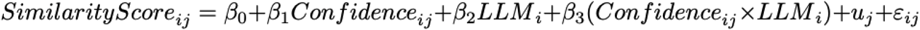

Where *β*_0_ is the fixed intercept, *β*_1_ represents the fixed effect coefficients, 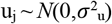 is the random intercept for dataset *j*, and *ε*_*ij*_ ∼ *N*(0,*σ*^2^) is the residual error for observation *i* in dataset *j*.

Post-hoc pairwise comparisons between LLMs used estimated marginal means (emmeans R package) with Tukey adjustment for multiple comparisons. Statistical significance was assessed at α = 0.05.

### Correlation Analysis

Relationships between confidence and similarity scores were analysed using Pearson correlation with degrees of freedom reported. Relationships between confidence ratings and heterogeneity were analysed using Spearman correlation due to ordinal confidence categories.

### Reproducibility Analysis

Standard deviation of similarity to author scores across multiple iterations of Immune Cell Atlas was computed per cluster per LLM. Consensus Cell Ontology terms were identified when all iterations produced identical terms. The proportion of clusters without consensus was calculated per LLM.

