## Supplementary Information 1 for "Multi-agent AI enables evidence-based cell annotation in single-cell transcriptomics"

### CyteOnto: Semantic Cell Type Annotation Comparison Using Large Language Models and Cell Ontology

The CyteOnto framework leverages the Cell Ontology (CL) and large language models (LLMs) to compute semantic similarity between cell type annotations. Our approach utilizes embedding-based semantic representations of LLM-generated cell type descriptions, both for computing the nearest mapping to a cell ontology term and for similarity between two cell ontology terms. Through systematic evaluation using three curated test datasets, we demonstrate that CyteOnto provides more biologically meaningful similarity scores compared to string-based and graph-based methods. We utilize the CyteOnto framework to evaluate CyteType against established cell type annotation tools, and to compare the performance of different LLMs within CyteType.

#### Semantic Similarity Computation Framework

CyteOnto implements a multi-step process to compute semantic similarity between cell type annotations. The framework consists of the following key steps:

1. **Cell Type Description Generation**: For each cell type annotation, we generate a detailed description using a LLM.
2. **Embedding Generation**: We convert the generated descriptions into high-dimensional vectors using text embedding models for semantic representation.
3. **Cell Ontology Mapping:** We map each cell type annotation to the nearest cell ontology term based on the cosine similarity of cell type description embeddings and precomputed embeddings of cell ontology terms.
4. **Similarity Computation**: We calculate the semantic similarity between two cell type annotations by determining a similarity score based on the mapped cell ontology terms.

#### Large Language Model Description Generation

CyteOnto generates a comprehensive textual description for each ontology term using LLMs such as Kimi-K2. The description is obtained by prompting the LLM with the cell type name and requesting a detailed description that includes the following key characteristics:

1. **Descriptive name**: A descriptive and concise name for the cell type.
2. **Functional characteristics**: Primary cellular functions and roles.
3. **Markers**: Marker genes that are commonly associated with the cell type.
4. **Disease relevance**: Associations with diseases or pathological conditions.
5. **Developmental stage**: Typical developmental stage or lineage of the cell type.

A specialized service that can retrieve up-to-date scientific abstracts from the NCBI PubMed database using a natural language query is also available to the description generation LLM as a tool. These characteristics are combined to form a comprehensive description for each cell type, for the generation of embeddings. For example, a cell type "Lymphatic Endothelial Cell" might yield the following description:

1. **Descriptive name**: Lymphatic vessel endothelial cell forming lymphatic vascular network throughout tissues.
2. **Functional characteristics**: Forms the inner lining of lymphatic vessels for lymph transport, fluid homeostasis, immune cell trafficking, and lipid absorption from the intestine.
3. **Markers**: PROX1, LYVE1, VEGFR3, PDPN, CCL21, ICAM1, VCAM1, NRP2
4. **Disease relevance**: Critical in lymphedema, cancer metastasis, inflammatory disorders, and lymphangiogenesis during tumor progression
5. **Developmental stage**: Differentiates from venous endothelial cells during embryonic development under PROX1 transcriptional control.

These characteristics are concatenated to form a comprehensive description as follows:

| |
|---|
| Lymphatic Endothelial Cell is a Lymphatic vessel endothelial cell forming lymphatic vascular |

network throughout tissues. Forms the inner lining of lymphatic vessels for lymph transport, fluid homeostasis, immune cell trafficking, and lipid absorption from the intestine. Critical in lymphedema, cancer metastasis, inflammatory disorders, and lymphangiogenesis during tumor progression. Differentiates from venous endothelial cells during embryonic development under PROX1 transcriptional control. The marker genes are PROX1, LYVE1, VEGFR3, PDPN, CCL21, ICAM1, VCAM1, NRP2.

#### Embedding-Based Semantic Representation

Generated descriptions are converted to high-dimensional vectors using state-of-the-art text embedding models (such as Qwen3-Embedding-8B). These embeddings capture the semantic meaning of the cell type descriptions and enable quantitative comparison across cell types. The CyteOnto package includes precomputed embeddings using the Qwen3-Embedding-8B model for the descriptions of all cell ontology terms generated using Kimi-K2.

#### Similarity Metric Design and Implementation

CyteOnto computes the similarity between two cell type annotations based on their mapped cell ontology terms. CyteOnto provides the following similarity metrics:

##### Set-based Similarity Measures

**Jaccard Similarity**: Standard set intersection over union using ancestor sets (including the terms themselves):

$$\text{Jaccard}(A_1, A_2) = \frac{|A_1 \cap A_2|}{|A_1 \cup A_2|}$$

**Set Cosine Similarity**: Normalized intersection based on geometric mean of set sizes:

$$\text{Set Cosine}(A_1, A_2) = \frac{|A_1 \cap A_2|}{\sqrt{|A_1| \cdot |A_2|}}$$

**Weighted Jaccard**: Incorporates ontological depth to emphasize more specific relationships:

$$\text{Weighted Jaccard} = \frac{\sum_{a \in A_1 \cup A_2} \min(w_d(a), w_d(a))}{\sum_{a \in A_1 \cup A_2} \max(w_d(a), w_d(a))}$$

Where $w_d(a) = \text{depth}(a)$ represents the ontological depth of the ancestor $a$, and $A_1, A_2$ represents the ancestor sets for terms 1 and 2, respectively.

#### Edge Weight-based Similarity Measures

**Weighted Ancestors Jaccard**: Uses ancestor count to weight relationships:

$$\text{Weighted Ancestors} = \frac{\sum_{a \in A_1 \cup A_2} \min(w_1^{anc}(a), w_2^{anc}(a))}{\sum_{a \in A_1 \cup A_2} \max(w_1^{anc}(a), w_2^{anc}(a))}$$

Where $w_i^{anc}(a) = \frac{1}{\max(|\text{ancestors}(a)|, 1)}$ if $a \in A_i$, else 0.

**Weighted Specificity Jaccard**: Emphasizes term importance using relative depth normalization:

$$\text{Weighted Specificity} = \frac{\sum_{a \in A_1 \cup A_2} \min(w_1^{spec}(a), w_2^{spec}(a))}{\sum_{a \in A_1 \cup A_2} \max(w_1^{spec}(a), w_2^{spec}(a))}$$

Where $w_i^{spec}(a) = \frac{\text{depth}(a)}{\max(\text{depth}(\text{term}_i), 1)}$ if $a \in A_i$, else 0.

**Weighted Cosine Jaccard**: Incorporates semantic similarity from embeddings as edge weights:

$$\text{Weighted Cosine} = \frac{\sum_{a \in A_1 \cup A_2} \min(w_1^{cos}(a), w_2^{cos}(a))}{\sum_{a \in A_1 \cup A_2} \max(w_1^{cos}(a), w_2^{cos}(a))}$$

Where $w_i^{cos}(a) = \text{cosine}(\mathbf{e}_a, \mathbf{e}_{\text{term}_i})$ if $a \in A_i$ and embeddings exist, else 0.

#### Path-based Similarity

**Path Similarity**: Measures similarity based on the shortest path through the lowest common ancestor (LCA):

$$\text{Path Similarity} = \frac{1}{\frac{d_1 + d_2}{2}}$$

Where $d_1 = \text{depth}(\text{term}_1) - \text{depth}(\text{LCA})$ and $d_2 = \text{depth}(\text{term}_2) - \text{depth}(\text{LCA})$.

#### Embedding-based Similarity

**Direct Cosine Similarity**: Standard cosine similarity between embedding vectors:

$$\text{Cosine}(\mathbf{v_1}, \mathbf{v_2}) = \frac{\mathbf{v_1} \cdot \mathbf{v_2}}{|\mathbf{v_1}||\mathbf{v_2}|}$$

**Ensemble Cosine Similarity**: Combines ontology matching confidence with embedding similarity:

$$\text{Ensemble Cosine} = \frac{s_1 + s_2 + \text{cosine}(\mathbf{v_1}, \mathbf{v_2})}{3 \cdot s_{\max}}$$

Where $s_1$ and $s_2$ are the embedding similarity scores between the input terms and their matched ontology terms, and $s_{\max}$ is the maximum pairwise similarity in the ontology embedding space.

**Gaussian Hill Kernel over Cosine (GHKcos)**: A kernel function designed to emphasize high-similarity relationships while maintaining sensitivity across the similarity spectrum:

$$\text{Gaussian Hill}(x) = A \cdot \exp\left(-\frac{(x-c)^2}{2\sigma^2}\right)$$

where $c$ is the center (default 1.0), $\sigma$ controls the width (optimally 0.25), $A$ is the amplitude (default 1.0), and $x$ is the raw cosine similarity score.

**Importance and Selection of the Default Kernel**

The application of a kernel function is critical for transforming raw cosine similarity scores into better discriminative measures. The Gaussian Hill Kernel reshapes the similarity distribution, increasing the contrast between highly similar pairs and less similar pairs. This non-linear transformation preserves the expected monotonic decay in similarity scores across a gradient, such as from a parent cell type to increasingly distant relatives. Through systematic parameter optimization, a kernel width of $\sigma = 0.25$ was identified as the optimal value. By default, CyteOnto uses this cosine similarity metric with the applied Gaussian Hill Kernel (GHKcos) to quantify semantic similarity.

**Kernel Parameter Optimization**

We systematically evaluated six Gaussian hill kernel widths ($\sigma$ = 0.15, 0.20, 0.25, 0.30, 0.35, 0.40) to determine optimal parameter settings. We computed cosine similarities between the Qwen3-Embedding-8B computed embedding vectors of the Kimi-K2-generated descriptions of cell ontology terms. Figure SI-

1-1 shows the effect of different kernel widths on original similarity scores. The optimization process considered:

- **Discrimination Power**: Ability to distinguish between biologically meaningful similarity levels.
- **Monotonicity Preservation**: Maintenance of expected ordering relationships.
- **Sensitivity Range**: Effective utilization of the full similarity spectrum.

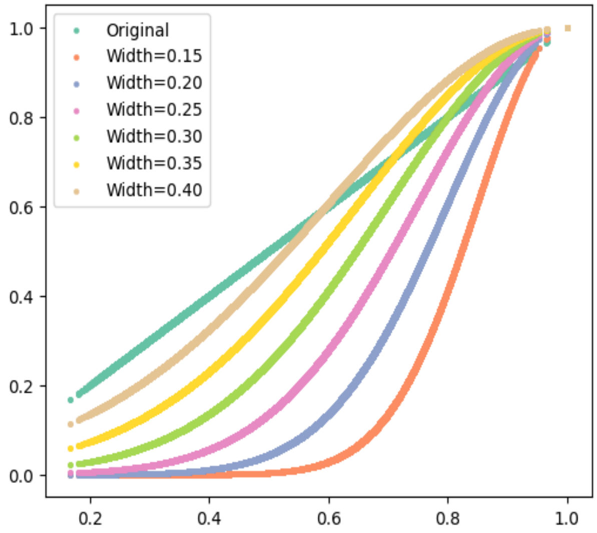


Figure SI-1-1: **Effect of the Gaussian Hill Kernel Widths on Pairwise Similarity Scores of Cell Ontology Terms**. A kernel width of σ = 0.25 was identified as the optimal value.

#### Evaluation Dataset Design and Validation

To evaluate the similarity metrics, we designed three carefully curated test datasets with distinct characteristics:

##### Dataset 1: Cell Ontology Term Pairs

Contains 30 pairs of CL identifiers with various biological relationships:

- **Identical terms**: Perfect similarity controls (CL:0000084 vs CL:0000084).
- **Parent-child relationships**: Direct hierarchical connections.
- **Sibling relationships**: Terms with shared parents but distinct functions.
- **Distant relationships**: Unrelated cell types from different lineages.

- **Specificity gradients**: Broad terms vs. highly specific subtypes.

##### Dataset 2: *CD4+, alpha-beta T cell* Reference Series

An evaluation series using *CD4+, alpha-beta T cell* (CL:0000624) as a fixed anchor against 40 progressively more distant cell types:

- **Identity match**: CD4+ T cell vs itself.
- **Direct children**: T helper subtypes (Th1, Th2, Treg).
- **Sibling T cells**: CD8+ T cells, memory/naive variants.
- **Related immune cells**: Other lymphocytes, dendritic cells, macrophages.
- **Hematopoietic relatives**: Erythrocytes, platelets, stem cells.
- **Distant cell types**: Neurons, hepatocytes, epithelial cells.

##### Dataset 3: *Sensory Neuron* Reference Series

Similar to the CD4+ series but focused on neuronal cell types, using the *sensory neuron* (CL:0000101) as an anchor against 40 cell types spanning neural and non-neural categories.

#### Results

Our evaluation compares the performance of the embedding-based GHKcos metric ($\sigma = 0.25$) with that of ontology-based methods across all three curated datasets.

##### Dataset 1: Cell Ontology Term Pairs

In this dataset, which evaluates a diverse range of pairwise relationships, the GHKcos ($\sigma = 0.25$) metric provided a stable and biologically plausible similarity curve. As seen in Figure SI-1-2A and Figure SI-1-2B, traditional metrics such as Jaccard, Set Cosine, and Path Similarity exhibit high volatility and non-monotonic behavior. For example, these methods often produce erratic scores that do not reflect known biological relatedness. In contrast, the embedding-based kernel metric exhibits a smoother profile that aligns more closely with qualitative biological understanding.

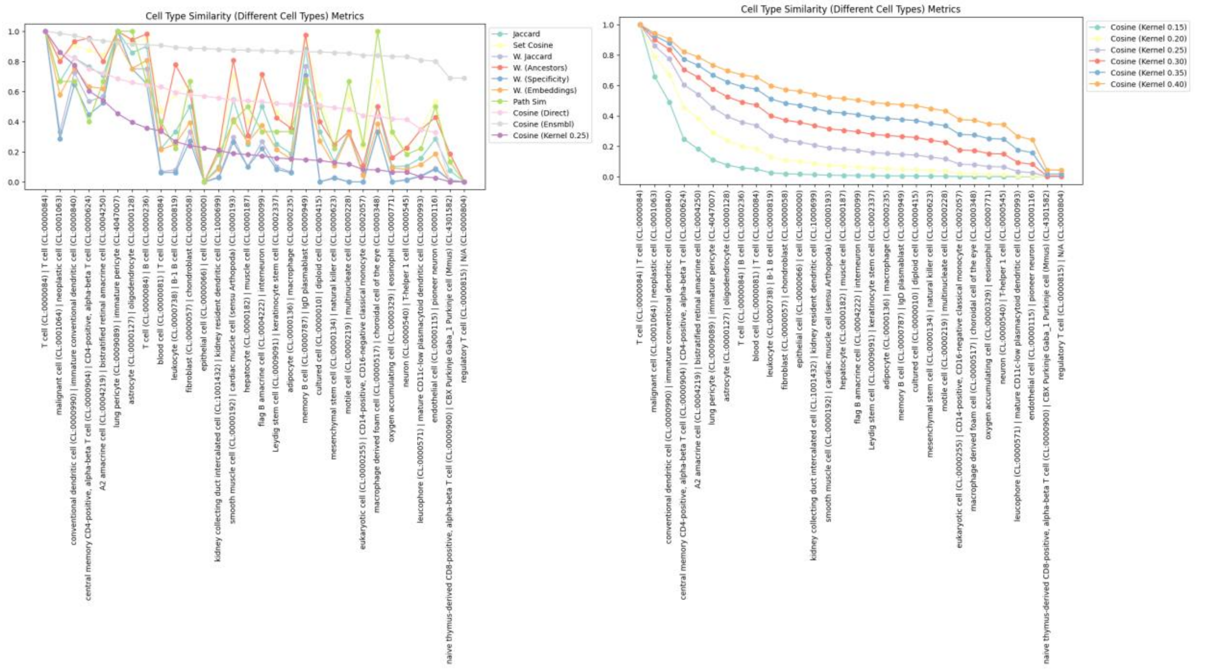


Figure SI-1-2: **Performance Comparison on the Cell Ontology Term Pairs Dataset**. **A)** Comparison of GHKcos (σ = 0.25) against other set-based, path-based, and embedding-based metrics. **B)** Comparison of the GHKcos metric across the six evaluated kernel widths.

**Dataset 2: CD4+ T Cell Reference Series**

The CD4+ T cell reference series provides a clear test of monotonicity. The GHKcos (σ = 0.25) metric excelled in this test, demonstrating a smooth, monotonically decreasing similarity score as the compared cell types become progressively more distant from the CD4+ T cell anchor. The similarity is highest for the identical pair, decreases slightly for its direct children (e.g., T-helper 1 cell) and siblings (e.g., CD8-positive, alpha-beta T cell), and continues to fall for more distant hematopoietic relatives before reaching a floor for non-related cell types like neurons and epithelial cells, as seen in Figure SI-1-3B. This behavior starkly contrasts with graph-based metrics, which, as shown in Figure SI-1-3A, produce highly erratic and noisy scores that fail to capture this clear biological gradient.

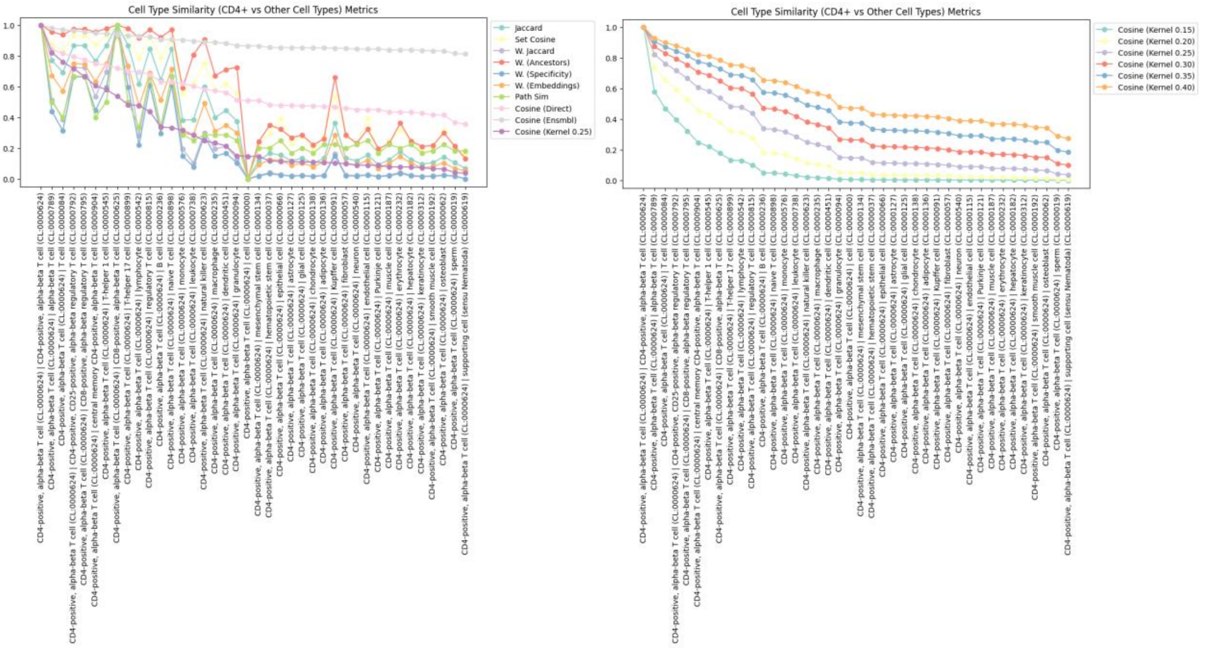


Figure SI-1-3: **Metric Evaluation on the CD4+ T Cell Reference Series**. **A)** Comparison of GHKcos (σ = 0.25) against other set-based, path-based, and embedding-based metrics. **B)** Comparison of the GHKcos metric across the six evaluated kernel widths.

**Dataset 3: Sensory Neuron Reference Series**

The results from the sensory neuron reference series are similar to the findings from the CD4+ T cell reference dataset. Once again, the GHKcos (σ = 0.25) metric produced a smooth, monotonic decay in similarity scores, starting high for closely related neurons (e.g., afferent neurons) and systematically decreasing for more distant cell types (Figure SI-1-4B) as compared to other metrics (Figure SI-1-4A).

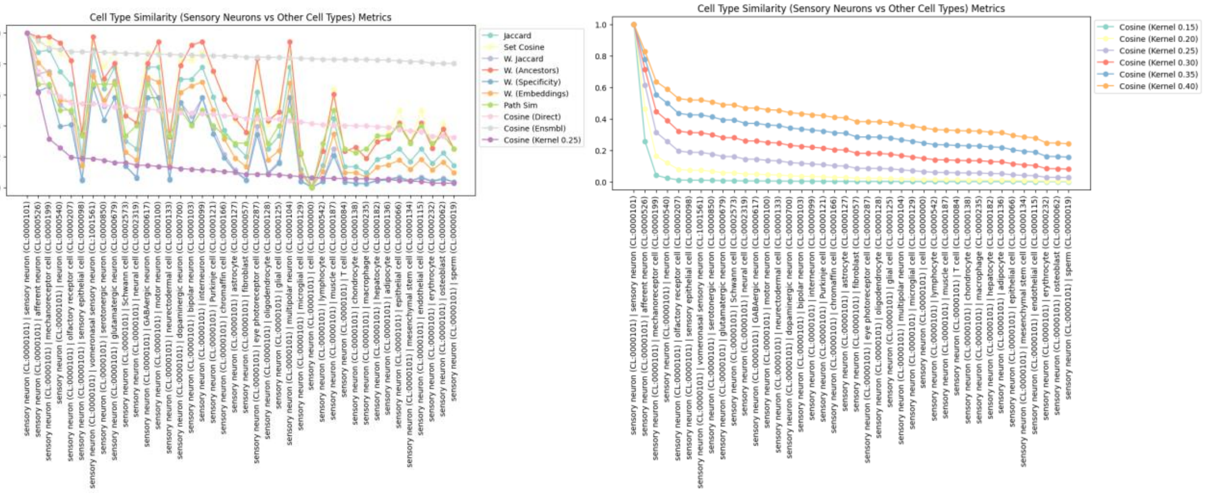


Figure SI-1-4: **Metric Validation on the Sensory Neuron Reference Series**. **A)** Comparison of GHKcos ($\sigma = 0.25$) against other set-based, path-based, and embedding-based metrics. **B)** Comparison of the GHKcos metric across the six evaluated kernel widths.
