## Supplementary Information 2 for "Multi-agent AI enables evidence-based cell annotation in single-cell transcriptomics"

### Benchmarking LLMs

#### 1. Supplementary Data

##### 1.1. Cost

Token usage for data was multiplied by the cost per token for both input and output, followed by the addition of the input and output costs. Cost varied dramatically, with open-weight LLMs often annotating clusters for less than a cent and some closed-weight models requiring greater than 25 cents to annotate a cluster (Table SI-2-1). In order to investigate the effect of the input data on cost, intraclass correlation coefficients (ICC) were calculated from a linear mixed effects model (LME). Variance decomposition showed costs were dictated by the choice of LLMs, with negligible variance attributable to the dataset ($ICC_{annotator}$ = 1.63%, $ICC_{reviewer}$ = 0.00%) and iteration ($ICC_{annotator}$ = 0.26%).

##### 1.2. Time

Likewise, the time required varied dramatically, with some LLMs requiring less than ten seconds to annotate a cluster (e.g., LLaMA Maverick 4) and others requiring over three minutes to do so (Table SI-2-1). The total runtime per dataset is dramatically reduced by parallelizing the analysis by cluster. In order to determine the effect of input data on runtime, ICCs were once again calculated from an LME. Here, minimal variance in run time was attributable to the dataset ($ICC_{annotator}$ = 5.46%, $ICC_{reviewer}$ = 1.98%) or iteration ($ICC_{annotator}$ = 2.51%).

##### 1.3. Tool Calling

Finally, to determine the effect of input data on tool calling, ICCs were calculated from an LME. Again, negligible variance in tool calling was explained by the dataset ($ICC_{annotator}$ = 0.33%, $ICC_{reviewer}$ = 0.97%) or iteration ($ICC_{annotator}$ = 0.01%).

**Table SI-2-1: Cost and Runtime per Large Language Model.** The cost and runtime per cluster of large language models to annotate and review. Values are estimated marginal means with a 95% confidence interval.

| **LLM** | **Cost per Cluster ($)** | | **Time per Cluster (Seconds)** | |
|---|---|---|---|---|
| | **Annotation** | **Reviewer** | **Annotator** | **Reviewer** |
| Claude Sonnet 4 | 0.15<br>[0.14, 0.15] | 0.14<br>[0.14, 0.15] | 96.04<br>[74.19, 117.89] | 139.78<br>[75.83 203.74] |
| DeepSeek R1 | 0.0093<br>[0.0051, 0.014] | 0.0078<br>[0.0027, 0.013] | 53.52<br>[31.67, 75.37] | 85.60<br>[21.64, 149.55] |
| Gemini 2.5 Flash | 0.012<br>[0.0073, 0.016] | 0.012<br>[0.0071, 0.017] | 20.68<br>[0.00, 42.53] | 45.30<br>[0.00, 109.25] |
| Gemini 2.5 Pro | 0.19<br>[0.18, 0.19] | 0.18<br>[0.18, 0.19] | 158.49<br>[136.63, 180.34] | 196.77<br>[132.82 260.73] |
| LLaMA 4 Maverick | 0.0039<br>[0.00, 0.0082] | 0.0027<br>[0.00, 0.0079] | 8.64<br>[0.00, 30.49] | 23.08<br>[0.00, 87.03] |
| Minimax M1 | 0.012<br>[0.0059, 0.019] | 0.024<br>[0.019, 0.030] | 315.12<br>[290.04, 340.20] | 867.07<br>[808.63, 925.50] |
| Magistral Medium | 0.048<br>[0.044, 0.052] | 0.046<br>[0.041, 0.051] | 23.32<br>[1.46, 45.17] | 48.27<br>[0.00, 112.23] |
| Kimi K2 | 0.011<br>[0.0069, 0.015] | 0.0097<br>[0.0046, 0.015] | 40.37<br>[18.52, 62.22] | 61.19<br>[0.00, 125.15] |
| GPT-4.1 | 0.058<br>[0.053, 0.062] | 0.054<br>[0.049, 0.059] | 48.58<br>[26.73, 70.43] | 82.61<br>[18.66, 146.57] |
| GPT-5 | 0.095<br>[0.091, 0.10] | 0.21<br>[0.20, 0.21] | 140.81<br>[118.76, 162.86] | 380.73<br>[317.56, 443.89] |
| GPT-oss 120B | 0.0042<br>[0.00, 0.019] | 0.0036<br>[0.00, 0.013] | 17.09<br>[0.00, 59.08] | 17.79<br>[0.00, 87.97] |
| Qwen3 235B A22B | 0.0033 | 0.0018 | 32.65 | 42.63 |

| | | | | |
|---|---|---|---|---|
| | [0.00, 0.0076] | [0.00, 0.0070] | [10.80, 54.50] | [0.00, 106.59] |
| Qwen3 235B A22B (Thinking) | 0.0057 [0.0013 0.010] | 0.0019 [0.00, 0.0070] | 204.19 [182.12, 226.25] | 127.60 [64.67, 190.52] |
| Qwen3 30B A3B (Thinking) | 0.0089 [0.00032, 0.018] | 0.0083 [0.00, 0.018] | 208.93 [180.14, 237.72] | 193.28 [123.09, 263.46] |
| Grok 4 | 0.28 [0.28, 0.29] | 0.26 [0.26, 0.27] | 311.30 [289.44, 333.17] | 337.19 [273.23, 401.14] |
| GLM 4.5 | 0.0059 [0.0016, 0.010] | 0.0068 [0.0016, 0.012] | 50.97 [29.12, 72.82] | 94.89 [30.93, 158.84] |

### 2. Accuracy

#### 2.1. Majority Agreement

In order to see whether LLMs significantly affected the majority agreement, the full generalised linear mixed effects model was compared to a null model. A likelihood ratio test comparing the two models showed majority class accuracy was significantly affected by LLMs ($X^2(15) = 67.99$, $p < .001$). When compared pairwise, Qwen3 235B A22B (both thinking and non-thinking variants), Grok 4, and LLaMA 4 Maverick all performed significantly worse than the leading LLM (Claude Sonnet 4) (Figure SI-2-1).

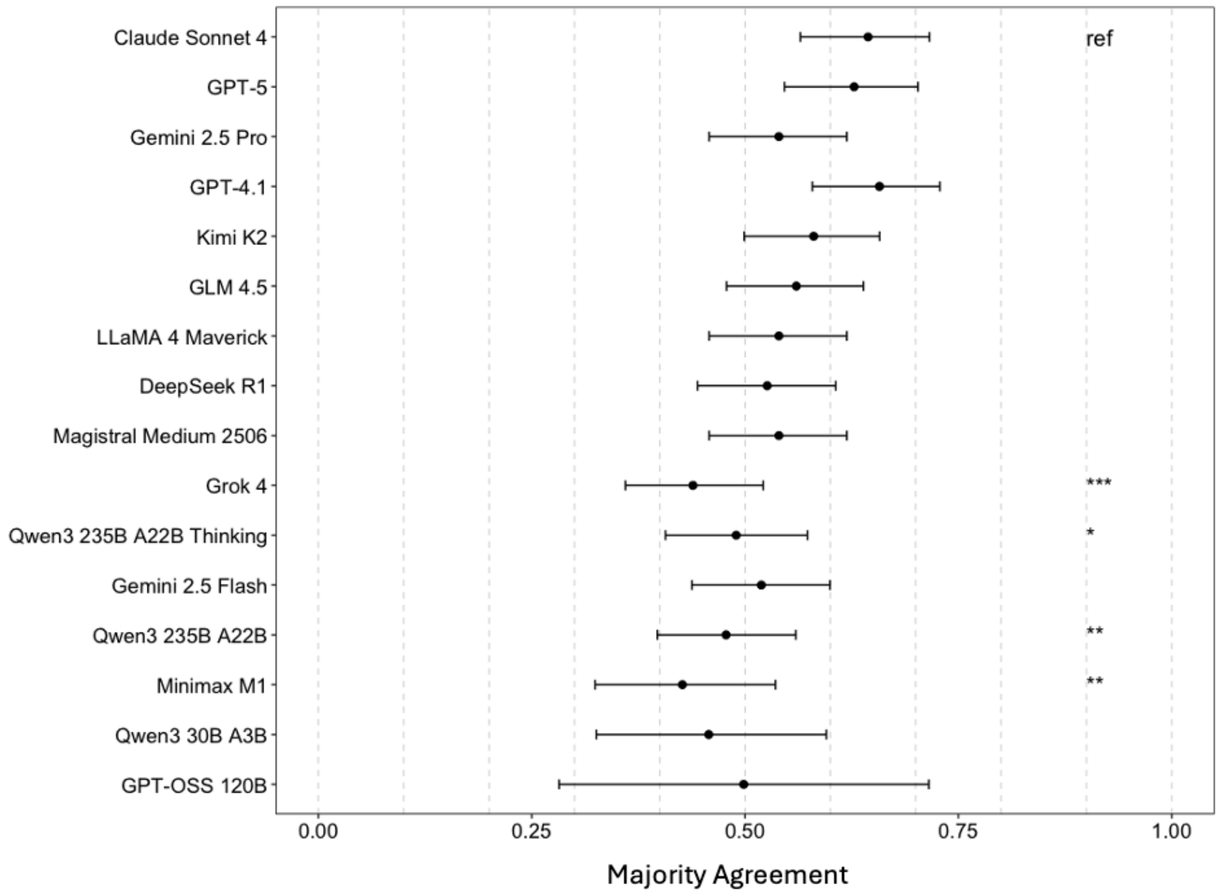


**Figure SI-2-1: Majority Agreement per Large Language Model.** The majority agreement of large language models annotating clusters. Values are estimated marginal means with 95% confidence interval; Tukey adjusted pairwise significance is referenced to leading LLM indicated by, with * indicating $p < .05$, ** indicating $p < .01$, and *** indicating $p < .001$.

#### 2.2. Similarity to Author

To see whether LLMs significantly affected the similarity to the author score, the full LME was compared to a null model. A likelihood ratio test comparing the two models showed that the similarity to author score was significantly affected by LLMs ($X^2(15) = 54.13$, $p < .001$). When compared pairwise, Qwen3 235B A22B (non-thinking) and Minimax M1 both performed significantly worse than the leading LLM (Gemini 2.5 Pro) (Figure SI-2-2).

To examine the relationship between model characteristics regression analysis was undertaken. Open-weight LLMs have a significantly reduced similarity to author score than their closed-weight counterparts ($b$ = -0.035, $SE$ = 0.011, $t(3977) = -3.33$, $p < .001$). Conversely, the increase in similarity score from reasoning was nonsignificant ($b$ = 0.014, $SE$ = 0.011, $t(3977)$, $p = .22$). Additionally, both graduate-level Google-proof Q&A diamond and massive multitask language understanding benchmarks were significant predictors of similarity to author score ($b$ = 0.0024, $SE$ = 0.00077, $t(3977) = 3.17$, $p < 0.01$ and $b$ = 0.0050, $SE$ = 0.0017, $t(3977) = 2.92$, $p < .001$, respectively) [1,2].

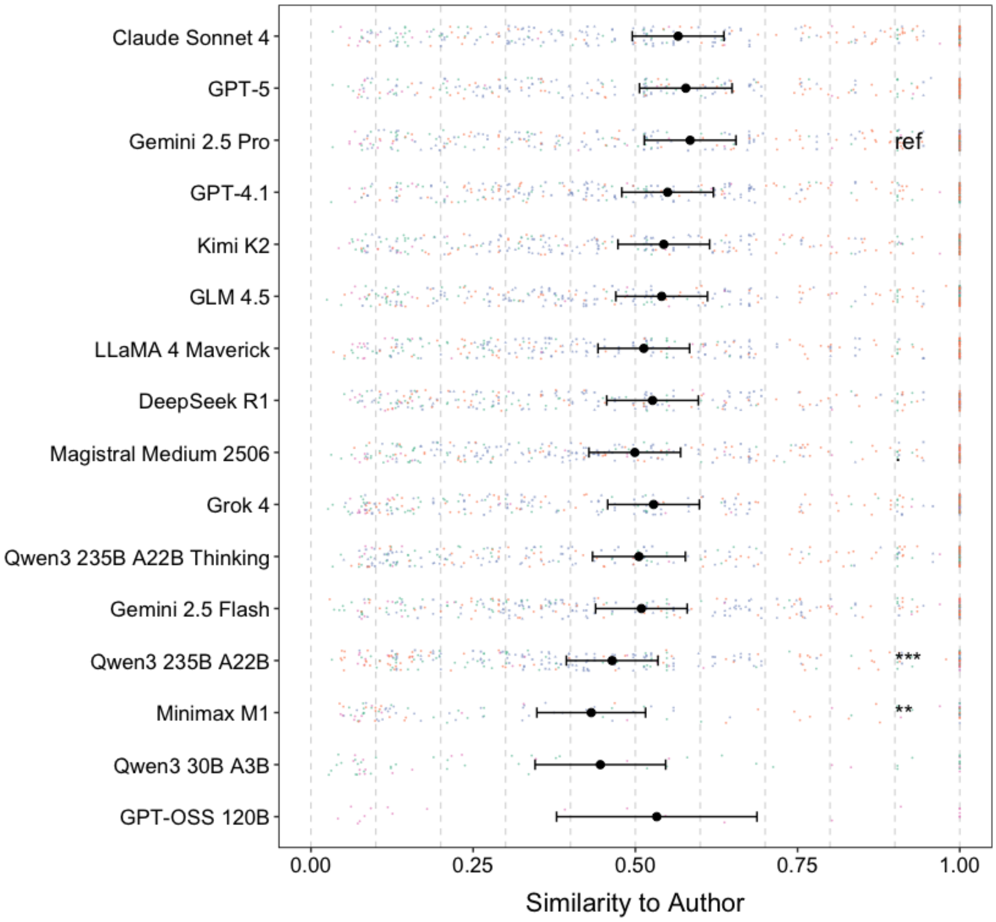


**Figure SI-2-2: Similarity to Author per Large Language Model.** The similarity to the author of large language models annotating clusters. Values (black) are estimated marginal means with 95% confidence interval, and jittered dots are raw values per cluster split by Human Immune Cell Atlas (sky blue), GTEx v9 (soft orange), HypoMap (seafoam green), and Murine Pancreatic Atlas (soft pink). Tukey adjusted pairwise significance is referenced to the leading LLM indicted ref, with ** indicating $p < .01$ and *** indicating $p < .001$.

### 3. Reliability

#### 3.1. Determinism of Output

To gauge the determinism of the output of LLMs, LLMs were run on the same data multiple times. The median standard deviation of LLMs across clusters was below 20% of the similarity score for all LLMs. Mean similarity to author score across iterations was not significantly correlated with standard deviation between iterations in any LLMs. Across all iterations, LLMs achieved a consensus cell type more than 70% of the time except for Qwen 3 235B A22B (thinking), DeepSeek R1, GPT-5, and Magistral Medium. To measure the influence of the framework on determinism of output, GPTCelltype and CyteType (both using GPT-5) were compared. Here, a Wilcoxon signed-rank test showed CyteType to be more robust in a non-significant manner ($V(45) = 360$, $p = .17$, $r = .35$).

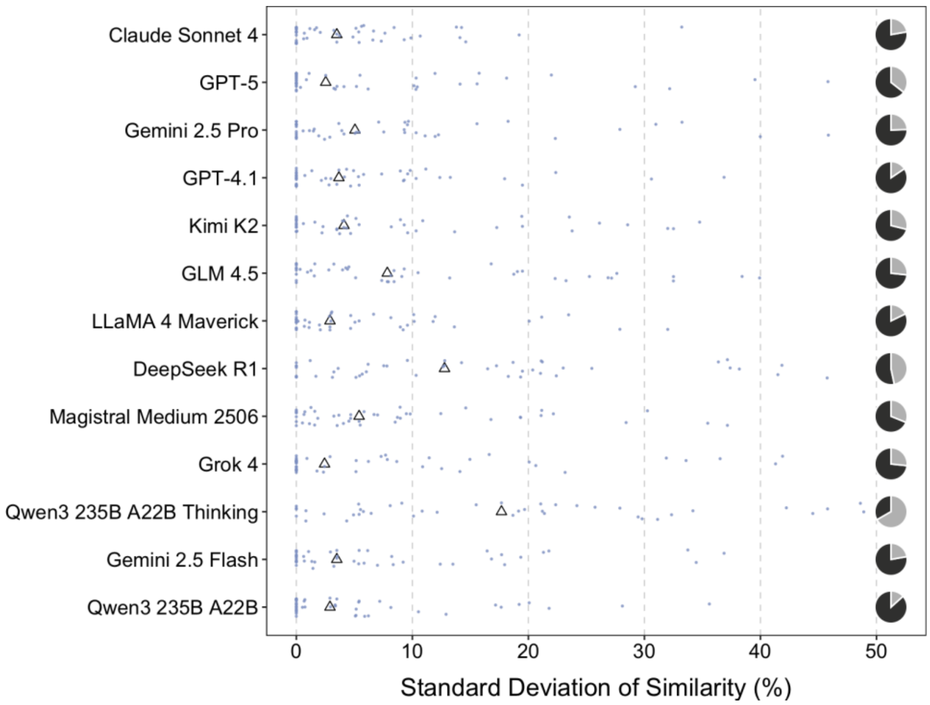


***Figure SI-2-3: Reliability per Large Language Model.*** The standard deviation of similarity to author scores across clusters for each LLM (%), with LLM medians (black triangles) and individual clusters (sky blue points).

The pie chart shows the proportion of clusters with a consensus cell ontology label across all iterations (black) versus those without consensus (grey).

#### 3.2. Heterogeneity Detection

LLMs within the reviewer agent analysed differential gene expression to predict whether clusters were heterogeneous. To verify the ability of CyteType to predict heterogeneous clusters, ROGUE was applied to cluster [3]. This orthogonal method produced little variance ($M$ = 0.889, $SD$ = 0.057), indicating most clusters were a pure population. An LME, accounting for dataset as a random effect, demonstrated CyteType's heterogeneity status significantly predicted ROGUE values ($b$ = -0.012, $SE$ = 0.0064, $t(3819) = -2.02$, $p$ = .043). To monitor the performance of LLM's heterogeneity ratings across multiple datasets an LME was utilised. Here, less heterogeneity was predictive of a greater similarity to the author score ($F(1, 3936) = 8.45$, $p < .01$). When accounting for the choice of LLMs, heterogeneity remained a significant predictor of similarity to the author score ($X^2(16) = 35.44$, $p < .01$).**SI-2**Heterogeneity ratings produced by GPT 5 were the most predictive of similarity to author score (Figure SI-2-4).

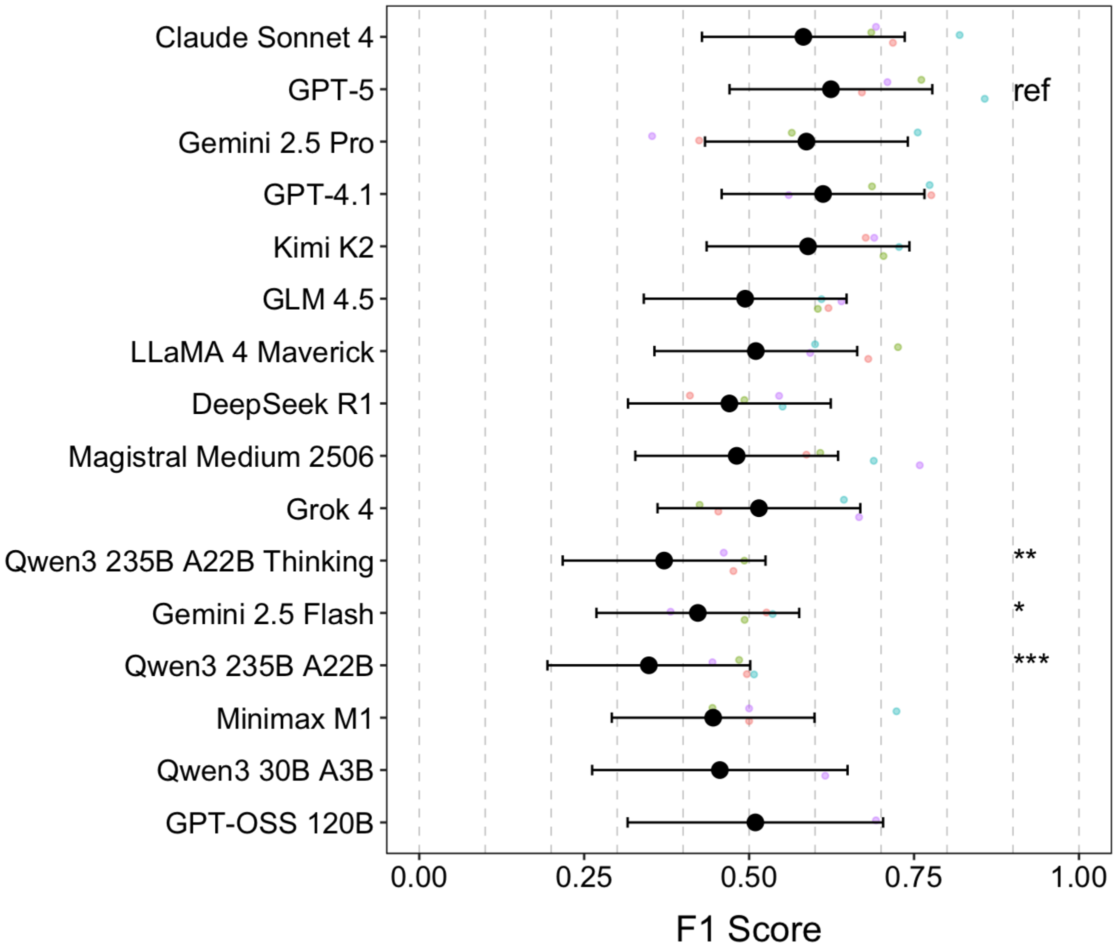


**Figure SI-2-4: Performance of Heterogeneity per Large Language Model.** The F1 of large language models in accurately predicting heterogeneous clusters. Values (black) are estimated marginal means with 95% confidence interval, and coloured dots are raw values per dataset; Tukey adjusted pairwise significance is referenced to the leading LLM indicated ref with * indicating $p < .05$ and ** indicating $p < .01$, and *** indicating $p < .001$.

#### 3.3. Reviewer Confidence

LLMs within the reviewer agent were exposed to the same set of annotations. These LLMs were then prompted to assign a confidence rating to the assigned cell type. Most LLM confidence ratings were moderate ($M$ = 0.61 [0.52, 0.71]), followed by high ($M$ = 0.31 [0.22, 0.39]), and low ($M$ = 0.11 [0.03, 0.19]) (Figure SI-2-5).

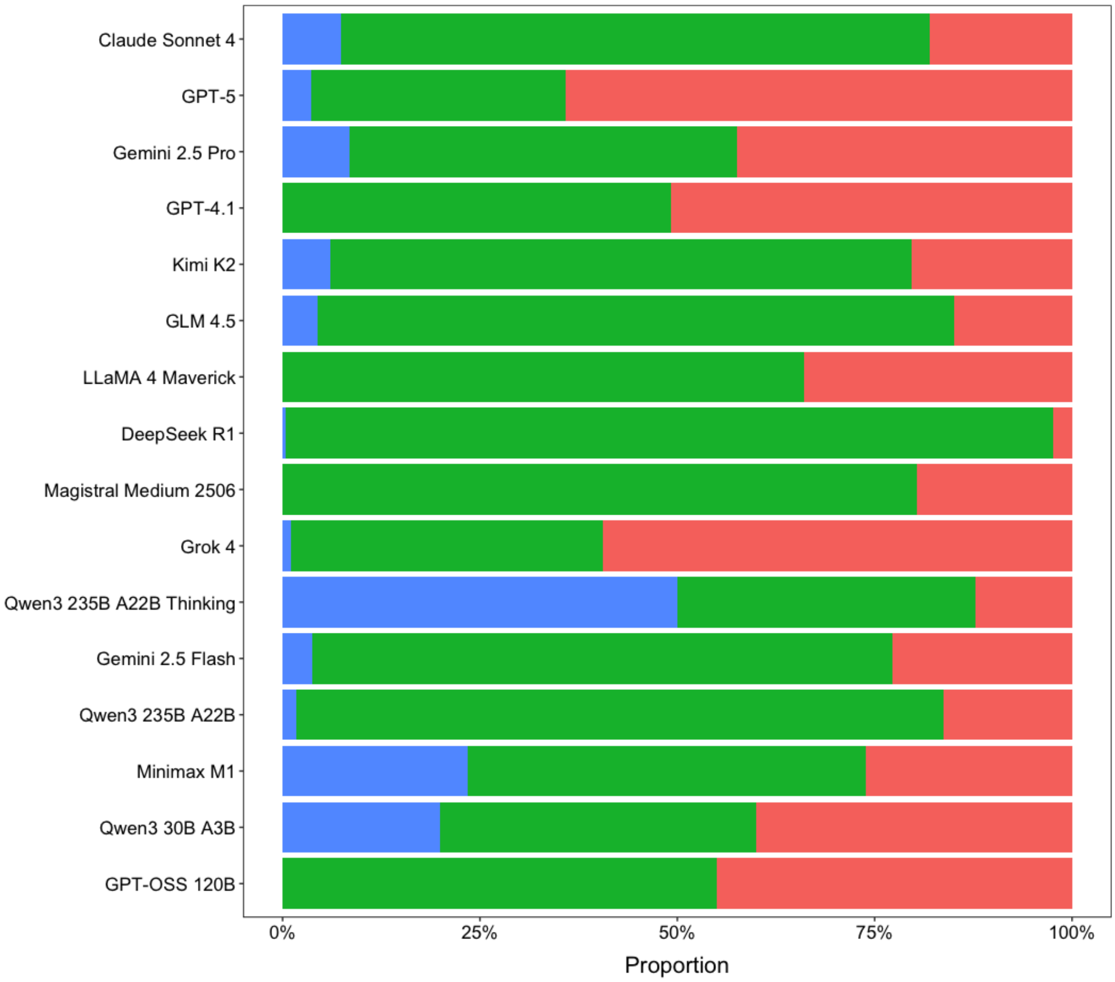


**Figure SI-2-5: Distribution of Confidence per Large Language Model.** The proportion of confidence rating of large language models in an annotation split by high (red), moderate (green), and low (blue).

Higher confidence was predictive of a greater similarity to the author score ($F(1, 3935.8) = 23.88$, $p < .001$); when accounting for the choice of LLMs, confidence remained a significant predictor of similarity to author score ($X^2(16) = 42.06$, $p < .001$). Estimated marginal means at discrete confidence levels (1 = Low, 2 = Moderate, and 3 = High) were utilised to create a post hoc regression of similarity scores across confidence levels (Table SI3-2). This indicated confidence was an important metric for self-assessment.

**Table SI-2-2: Relationship of Confidence to Similarity to Author per Large Language Model.**

Gradients of estimated marginal means from a linear mixed effects model relating confidence to similarity to the author. Values represent gradients with 95% confidence intervals.

| **LLM** | **Gradient [95% CI]** |
|---|---|
| Claude Sonnet 4 | 0.09 [0.01, 0.16] |
| DeepSeek R1 | 0.16 [-0.06, 0.38] |
| Gemini 2.5 Flash | -0.01 [-0.09, 0.06] |
| Gemini 2.5 Pro | 0.09 [0.03, 0.15] |
| LLaMA 4 Maverick | 0.00 [-0.08, 0.08] |
| Minimax M1 | 0.00 [-0.09, 0.08] |
| Magistral Medium | 0.08 [-0.01, 0.17] |
| Kimi K2 | -0.01 [-0.08, 0.06] |
| GPT-4.1 | 0.11 [0.04, 0.18] |
| GPT-5 | 0.00 [-0.07, 0.06] |
| GPT-oss 120B | 0.02 [-0.26, 0.30] |
| Qwen3 235B A22B | 0.07 [-0.02, 0.16] |
| Qwen3 235B A22B (Thinking) | 0.00 [-0.06, 0.05] |
| Qwen3 30B A3B (Thinking) | -0.11 [-0.29, 0.08] |
| Grok 4 | 0.08 [0.01, 0.15] |
| GLM 4.5 | 0.10 [0.02, 0.19] |
