## Supplementary Information 3 for "Multi-agent AI enables evidence-based cell annotation in single-cell transcriptomics"

### CyteType Reports

#### Benchmarked Datasets

**Table SI-3-1: Datasets Included in Benchmark** with down sampling notebook, as well as dataset and CyteType report.

| **Dataset** | **Colab Notebook** | **CyteType Report** | **Down sampled H5ad** |
|---|---|---|---|
| Tabula Sapiens | Link | Link | Link |
| Immune Cell Atlas | Link | Link | Link |
| GTEX v9 | Link | Link | Link |
| Mouse Pancreatic Cell Atlas | Link | Link | Link |
| HypoMap | Link | Link | Link |
| Diabetic Kidney Disease | Link | Link | Link |
| Human Lung Cell Atlas (Core) | Link | Link | Link |
| Human Cell Atlas | Link | Link | Link |
| CellHint Blood | | Link | |
| CellHint Bone Marrow | | Link | |
| CellHint Heart | | Link | |
| CellHint Hippocampus | | Link | |
| CellHint Intestine | | Link | |
| CellHint Kidney | | Link | |
| CellHint Liver | | Link | |
| CellHint Lung | | Link | |
| CellHint Lymph Node | | Link | |
| CellHint Pancreas | | Link | |
| CellHint Skeletal Muscle | | Link | |
| CellHint Spleen | | Link | |

#### CyteType Review

CyteType was applied to 20 datasets, with CyteOnto then applied to the resulting predictions. 41 % of author-denominated clusters were confirmed with additional enhancement (*e.g.*, additional information on cell state), 29 % were functionally refined, whilst 30% were majorly corrected (Table SI-4-2).

In order to see which cell types benefited the most from CyteType, individual cluster annotations were inspected. 56% of clusters in the Mouse Pancreatic Cell Atlas dataset clusters with low similarity to the author were predicted with high confidence by CyteType, indicating the most dramatic difference between authors and CyteType's predictions.

Cells mapped to cell ontology terms were compared groupwise to examine whether specific cell types were vulnerable to CyteType major reannotation. Across the major cell types examined (e.g., hematopoietic, neuronal, stem, contractile, and pancreatic), the fraction of clusters that required major reannotation was broadly consistent.

**Table SI-3-2: CyteOnto Evaluation of CyteType.** The percentage of clusters per dataset was highly, moderately, or lowly similar to the author.

| **Similarity to the Author** | **Percentage of Clusters per Dataset (%) [95% CI]** | **Impact** |
|---|---|---|
| High (66.6% - 100%) | 41 [34, 49] | Confirmed with enhancement |
| Moderate (33.3 - 66.6%) | 29 [24, 34] | Functional refinement |
| Low (0 - 33.3%) | 30 [21, 38] | Major corrections |

Predicted cell types were classified into subtypes utilising the hierarchical graph structure of cell ontology (e.g., haematological, neuronal, contractile). Additionally, a pancreatic subtype was created by the presence of "pancreatic" within cell ontology terms. Predictions were surveyed for the presence of cell types from these subtypes, in order to validate CyteType predictions.

Within all datasets, 972 different annotations were made, mapping to 327 unique cell ontology terms; no cell ontology term appeared more than 2.5% of the time (Figure SI-4-1-a). Additionally, 116 unique cell states were found, with activated (37%) and mature (16%) cell states being the most prevalent (Figure SI-4-1-b). CyteType was generally highly confident in its predictions ($M$=0.71 [0.65, 0.77]), except for the two pancreas datasets (Figure SI-4-1-c). CyteType had moderate confidence for 70% of the clusters in the Mouse Pancreatic Cell Atlas and 57% in the CellHint Pancreas dataset. CyteType predicted that more than half of clusters were heterogeneous ($M$=0.54 [0.48, 0.6]). The aforementioned pancreatic datasets both had greater than 75% of clusters predicted as heterogeneous. Confidence in predictions negatively correlated with predictions of heterogeneity ($r(18) = -0.79$, *95% CI* [-0.91, -0.53], $p < .001$).

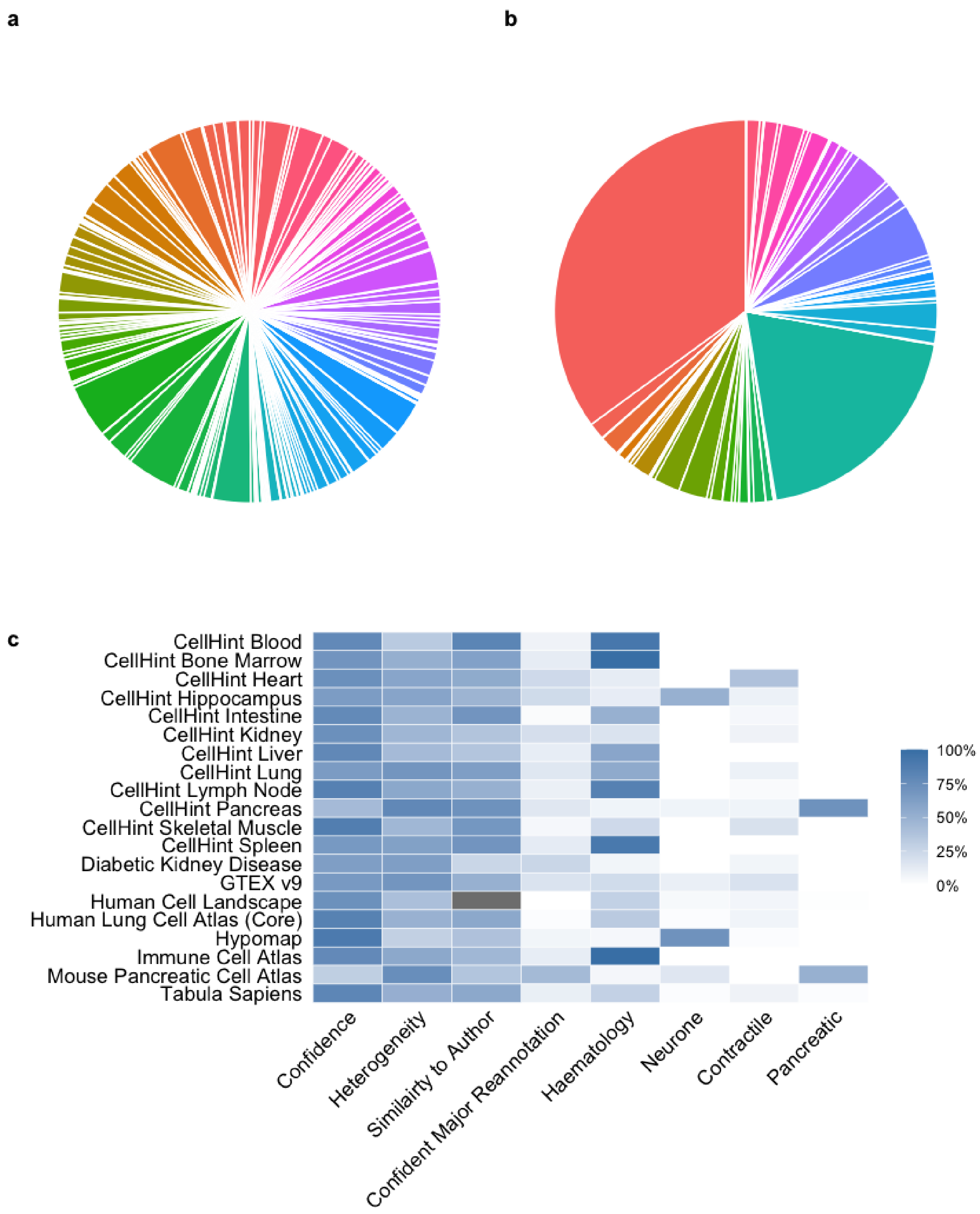


**Figure SI-3-1: CyteType Review in Extensive Benchmarking.** Pie charts with the proportion of unique cell ontology terms (a) and unique cell states (b). Heatmap showing proportion of clusters belonging to each category by dataset (c).
